# Evidence for the role of extrusomes in evading attack by the host immune system in a scuticociliate parasite

**DOI:** 10.1101/607622

**Authors:** Iria Folgueira, Jesús Lamas, Ana Paula De Felipe, Rosa Ana Sueiro, José Manuel Leiro

**Affiliations:** Departamento de Microbiología y Parasitología, Instituto de Investigación y Análisis Alimentarios, Campus Vida, Universidad de Santiago de Compostela, Spain; Departamento de Biología Funcional, Instituto de Acuicultura, Campus Vida, Universidad de Santiago de Compostela, Spain

**Keywords:** *Philasterides dicentrarchi*, turbot, exocytosis, extrusomes, trichocyst matrix proteins, mucin-like glycoproteins

## Abstract

Like other ciliates, the scuticociliate parasite of turbot, *Philasterides dicentrarchi*, produces only a feeding or growing stage called a trophont during its life cycle. Exposure of the trophonts to immune serum extracted from the host and containing specific antibodies that induce agglutination / immobilization leads to the production of a mucoid capsule from which the trophonts later emerge. We investigated how these capsules are generated, observing that the mechanism was associated with the process of exocytosis involved in the release of a matrix material from the extrusomes. The extruded material contains mucin-like glycoproteins that are deposited on the surface of the cell and whose expression increases with time of exposure to the turbot antibodies, at both protein expression and gene expression levels. Stimulation of the trophonts with host immune serum also causes an increase in discharge of the intracellular storage compartments of calcium necessary for the exocytosis processes in the extrusomes. The results obtained suggest that *P. dicentrarchi* uses the extrusion mechanism to generate a physical barrier protecting the ciliate from attack by soluble factors of the host immune system. Data on the proteins involved and the potential development of molecules that interfere with this exocytic process could contribute to the development of glycosylated recombinant vaccines and drugs to improve the prevention and control of scuticociliatosis in turbot.

## Introduction

Exocytosis may be an important mechanism of communication between microbes. Indeed, some microorganisms can develop highly specialized exocytotic organelles via the extrusion of different materials with important roles in mechanisms enabling adaptation to different environmental conditions [1]. Several groups of protozoa possess different types of exocytotic extrusive organelles, known as extrusomes. These organelles are associated with the cell membrane and have different structures containing a material that is usually expelled or extruded from the cell and that participates in different functions [2]. In ciliates, most extrusomes belong to the trichocyst type, which are characteristically spindle shaped and can quickly download their protein content in the form of a projectile in response to mechanical or physical stimuli, and with a probable function in defence against predators [3]. Other common extrusomes in some groups of ciliates include the toxicysts and haptocysts, which contain toxic material or can extrude material capable of penetrating the prey; both have a possible predatory function for prey capture and food uptake [4,5]. The function of trichites-type extrusomes, rod-shaped organelles circumferentially arranged in plasma pockets [6], is not yet completely known. However, it is believed that they can act as defensive or offensive elements [7]. Mucocysts and cortical granules, a special type of mucocysts, secrete an amorphous mucilaginous protective material on the cell surface. In some species, this material may be involved in the formation of cysts or temporary capsules with a protective role and constituting a first line of defence against predators in the ciliate, regulating cell ionic concentration and anchoring cells to substrates [3, 8-10].

*Philasterides dicentrarchi* is an amphizoic scuticociliate, originally free-living, which under certain conditions can be transformed into an opportunistic histiophagous parasite in cultivated flat fish, causing a serious disease called scuticociliatosis and producing high mortality rates [11,12]. In order to produce the parasitic phase, the ciliate must develop various strategies of biochemical adaptation to its new habitat [13,14]. In addition, it must evade attack by the fish immune system, especially by lysis induced by soluble factors in the serum, such as complement. Activation of complement via the classical route (in conjunction with antibodies), together with activation of the coagulation system, causes destruction of the parasite [15–17]. Two types of extrusomes have been characterized in *P. dicentrarchi*: one fusiform, compatible with trichocysts, and the other spherical, compatible with mucocysts, and which release a thin layer of mucus on the cell surface [18,19]. In previous studies, we have observed that incubation (for 2h) of *P. dicentrarchi* trophonts with serum from turbot that had survived a natural outbreak of scuticociliatosis caused agglutination and immobilization of the ciliates and the appearance of numerous capsules from which the trophonts later emerged. We interpreted this phenomenon as a possible antigenic change and a mechanism of evasion of the humoral immune response [20].

In the present study, we aimed i) to elucidate the role of the extrusomes in capsule production induced by incubation of the trophonts of *P. dicentrarchi* with immune sera from vaccinated turbot that produce agglutinating and immobilizing antibodies, ii) to characterize the proteins of the trichocysts and mucocysts after extrusion, and iii) to demonstrate the role of the process of exocytosis as a ciliate defence mechanism against attack by the soluble factors of the host humoral immune system.

## Materials and Methods

### Parasites

Specimens of *P. dicentrarchi* (isolate I1) were collected under aseptic conditions from peritoneal fluid obtained from experimentally infected turbot (*Scophthalmus maximus)*, as previously described [21]. The ciliates were cultured at 21 °C in complete sterile L-15 medium, as previously described [20]. In order to maintain the virulence of the ciliates, fish were experimentally infected every 6 months by intraperitoneal (ip) injection of 200 μL of sterile physiological saline containing 5×10^5^ trophonts, and the ciliates were recovered from ascitic fluid and maintained in culture as described above.

### Experimental animals

Turbot of approximately 50 g body weight were obtained from a local fish farm. The fish were held in 250-L tanks with aerated recirculating sea water maintained at 14 °C. They were subjected to a photoperiod of 12L:12D and fed daily with commercial pellets (Skretting, Burgos, Spain). The fish were acclimatized to laboratory conditions for 2 weeks before the start of the experiments.

Swiss ICR (CD-1) mice (eight to ten weeks old), supplied by Charles River Laboratories (USA), were bred and maintained in the Central Animal Facility of the University of Santiago de Compostela (Spain). The mice were reared following the criteria for the protection, control, care and welfare of animals and the legislative requirements relating to the use of animals for experimentation (EU Directive 86/609/EEC), the Declaration of Helsinki, and/or the Guide for the Care and Use of Laboratory Animals as adopted and promulgated by the US National Institutes of Health (NIH Publication No. 85–23, revised 1996). The Institutional Animal Care and Use Committee of the University of Santiago de Compostela approved all experimental protocols.

### Microscopic analysis

#### Scanning electron microscopy (SEM)

Ciliates treated with turbot immune serum (see Immunization and serum collection), were collected by centrifugation at 1000 × g, were fixed with 2.5% (v/w) glutaraldehyde in a cold solution of 4% paraformaldehyde in 0.1 M potassium phosphate buffer (PB), pH 7.2 for 30 min. The samples were post-fixed for 30 minutes with 1% (wt/v) osmium tetroxide in PB. The samples were then washed three times with distilled water and dehydrated in a series of ethanol (50, 70, 90, 95, 100, 100% for 10 min each) and hexamethyldisilazane (HMDS, Sigma-Aldrich) (50 and 100% for 10 min each). Finally, the samples were mounted on aluminium stubs, sputter coated with a layer of iridium, by using a Q150T-S sputter coater (Quorum Technologies, UK), and viewed under a Zeiss Fesem ultra plus microscope (Zeiss, Germany) at 10 kV.

#### Transmission electron microscopy (TEM)

For TEM, we followed the technique described by [19]. Briefly, the cultured ciliates were collected by centrifugation at 1000 × g for 5 min. Cells were fixed in 2.5% (v/v) glutaraldehyde in 0.1 M cacodylate buffer at pH 7.2. They were then washed several times with 0.1 M cacodylate buffer and post-fixed in 1% (wt/v) OsO_4_, pre-stained in saturated aqueous uranyl acetate, dehydrated through a graded acetone series and embedded in Spurr’s resin. Semi-thin sections were then cut with an ultratome (Leica Ultracut UCT, Leica microsystems, Germany) and stained with 1% toluidine blue for examination under a light microscope. Ultrathin sections were stained in alcoholic uranyl acetate and lead citrate and viewed in a Jeol JEM-1011 transmission electron microscope (Jeol, Japan) at an accelerating voltage of 100 kV.

#### Histochemistry: Safranin-O Staining

For detection of mucin-type proteins, the cells were stained with Safranin-O. Ciliates were incubated without turbot immune serum or with the serum for different times. The ciliates were fixed in 10% buffered formalin (PBS; 0.01 M Na_2_HPO_4_, 0.0018 M KH_2_PO_4_, 0.0027 M KCl, 0.137 M NaCl, pH 7.0). The samples were then washed 2 times with distilled water and incubated for 5 min with an aqueous solution of 0.1% of Safranin-O. After exhaustive washing with water to eliminate excess dye, the preparation was air-dried and mounted using a permanent mounting medium (Entellan^®^, Merck).

### Trichocyst associated proteins

The sequences of several mRNAs that encode proteins potentially related to the trichocysts of *Philasterides dicentrarchi* were obtained from a previous RNAseq study carried out to compare the transcriptome of several *P. dicentrarchi* strains, in collaboration with ZF-Screen (Holland). The assembled sequences were analyzed using Blastgo software 5.0 (Biobam, Spain), to identify homologous sequences, before functional annotation. Annotated sequences that encode proteins potentially related to the trichocysts of ciliates were selected using the BLASTx tool of the TGD Wiki (http://www.ciliate.org/blast/blast_link.cgi) where the *Tetrahymena thermophila* gene and protein sequences database is located. To confirm the nucleotide sequences that encode the proteins associated with the extrusomes obtained by RNAseq, their cDNAs were amplified by RT-PCR and sequenced by Sanger Sequencing (Eurofins Genomics, Germany). The selected proteins associated with the extrusomes were the *P. dicentrarchi* trichocyst matrix protein T2A (TMPT2A) (GenBank accession number MH412657.1) and the *P. dicentrarchi* trichocyst matrix protein T4-B (TMPT4B) (GenBank accession number MH412658.1).

### Production of recombinant proteins in yeast cells

The complete nucleotide sequence that encoded TMPT2A was modified and optimized to produce the recombinant protein in the yeast *Klyuveromyces lactis*, by using the bioinformatics tool developed by Integrated DNA Technology (IDT) (https://eu.idtdna.com/CodonOpt). The gene was then synthethized by Invitro GeneArt Gene Synthesis (ThermoFisher Scientific). For expression of recombinant protein in yeast, the *K. Lactis* Protein Expression kit (New England Biolabs, UK) was used with the pKLAC2 vector, following the instructions provided by the manufacturer. Initially, the synthesized nucleotide sequence was cloned in the pSpark^**®**^ II vector (Canvax, Spain), and the recombinant plasmid was subsequently amplified in competent *Escherichia coli* strain DH-5α. After extraction and purification of the plasmid from the bacteria, PCR was carried out using the following primers: FT2AKl 5’ CGCCTCGAGAAAAGAatgcgtgtctgaccgcacta-3’ / RT2AKl 5-’ ATAAGAATGCGGCCGCTTAATGATGATGGTGATGGTGATGATGGTGATGatcggcacgctttacgtc ga-3’. The reverse primer includes 10 codons encoding histidine at the C-terminal end of the protein. The yeasts were then transformed with the cloned pKLAC2 plasmid and seeded in YCB agar medium plates containing 5 mM acetamide at 30°C for 3–4 days until colony formation. Several of the colonies were collected and inoculated in the YPGal medium at 30 °C for 3-4 days with shaking at 250 rpm. When a suitable cell density was reached, the medium was centrifuged at 6000 xg for 10 min, and the supernatant was held at 4 °C until use. The protein was purified by submitting the supernatant to immobilized metal affinity chromatography (IMAC) using prepacked columns with Ni-Sepharose (HisTrap^TM^, GE Healthcare) in an ÄKTA Star protein purification system (GE Healthcare) following the manufacturer’s instructions. Once eluted, the protein was fully dialysed against distilled water using dialysis tubing of pore size 3 kDa. Finally, the protein was lyophilized and stored at 4 ° C until use.

### Sodium dodecyl sulphate polyacrylamide gel electrophoresis (SDS-PAGE)

SDS-PAGE analysis of the recombinant TMPT2A (rTMPT2A) was performed on linear 12.5% polyacrylamide minigels in a Mini-Protean^®^ Tetra cell system (BioRad, USA), as described by [22]. The gels consisted of 4% stacking gel and a 12.5% linear separating gel. Samples were dissolved in 62 mM Tris-HCl buffer (pH 6.8) containing 2% SDS, 10% glycerol and 0.004% bromophenol blue and heated for 5 min in a boiling water bath. The gels were electrophoresed at a constant 200 V in Tris-glycine electrode buffer (25 mM Tris, 190 mM glycine; pH 8.3). The gels were then fixed in 12% trichloroacetic acid for 1h and stained with QC Colloidal Coomassie stain (BioRad). Molecular weights were estimated using a calibration curve (Log_10_ MW vs Rf) constructed with a prestained protein standard (NZY Colour Protein Marker II, Nzytech, Portugal).

### Immunization and serum collection

Turbot were immunized by intraperitoneal injection (i.p.) on days 0 and 30 with 200 μl of an emulsion containing 10^6^ ciliates/mL inactivated with 0.2% formalin and 50% adjuvant Montanide ISA 763A (Seppic, France) [23]. Blood samples, obtained by caudal vein puncture, were allowed to clot for 2 h at room temperature before being centrifuged at 2000 x*g*. The serum was collected and stored at −20°C until use.

A group of five ICR (Swiss) CD-1 mice were immunized by i.p. injection with 200 μL per mouse of a 1:1 (v/v) mixture of Freund’s complete adjuvant (Sigma-Aldrich) and a solution containing 250 μg of purified rTMPT2A. The same dose of purified protein was prepared in Freund’s incomplete adjuvant and injected i.p. in mice 15 and 30 days after the first immunization. The mice were bled via retrobulbar venous plexus 15 days after the last immunization (Day 30) for initial checking of the antibody levels. If the antibody levels were satisfactory, the mice were decapitated and immediately bled. The blood was allowed to coagulate overnight at 4°C, and the serum was then separated by centrifugation (2000 xg for 10 min), mixed 1:1 with glycerol and stored at −20°C until use.

### Immunological assays

#### Immobilization/agglutination assay

Cultured ciliates were washed 3 times in incomplete L-15 medium. Aliquots of 200 ciliates were added to individual wells of 96 well microplates (Corning, USA), in a final volume of 50 μL in L5-medium. Before the assay, the serum was heat-inactivated at 56 °C for 30 min. The immune serum was assayed in triplicate and added to the wells containing the ciliates at dilutions of 1/25, 1/50 and 1/100 in L-15 medium. The plates were incubated at room temperature and checked for immobilization/agglutination responses, after 15, 30 and 60 min, under an inverted microscope (Nikon Eclipse TE300 Nikon, Japan). All assays included a ciliate control in incomplete L-15 medium with no serum. The agglutination response was expressed as the percentage of agglutinated ciliates.

#### Immunofluorescence and confocal microscopy

For immunolocalization of mucin-like proteins, an immunofluorescence assay was performed as previously described [24}. Briefly, 5×10^6^ ciliates incubated for different times with the immune serum from turbot, were centrifuged at 1000 xg for 5 min, washed twice with PBS pH 7.0 and fixed for 15 min in a solution of 4% formaldehyde in PBS at room temperature. The ciliates were then washed twice with PBS, resuspended in a solution containing 0⋅3% Triton X-100 in PBS for 3 min, washed twice with PBS, and incubated with 1% BSA for 30 min. After this blocking step, the ciliates were washed in PBS and incubated at room temperature with agitation (750 rpm) for an hour with a 1:100 dilution in PBS of mice serum anti-rTMPT2A. After being washed 3 times with PBS, the ciliate samples were added to a 1:1000 dilution of fluorescein isothiocyanate (FITC) conjugated rabbit/ anti-mouse Ig (DAKO, Denmark) and incubated for 1h at room temperature, in darkness. After another three washes in PBS, the samples were mounted in PBS-glycerol (1:1) and visualized by confocal microscopy (Leica TCS-SP2, Leica Microsystems, Germany).

#### Fluorescent enzyme-linked immunosorbent assay (FELISA)

For quantification of the expression of the TMPT2A by the trophonts incubated with turbot immune serum for 30 min and 6h, a FELISA was conducted as previously described [14]. Ciliate lysate (CL), prepared as previously described [20], was used as antigen in the assay. One μg of CL isolated from trophonts dissolved in 100 μL of carbonate-bicarbonate buffer pH 9.6, was added to wells of ELISA microplates (high binding, Greiner Bio-One, Germany) and incubated overnight at 4°C. The wells were then washed three times with 50 mM Tris, 0.15 M NaCl, pH 7.4 buffer (TBS), blocked for 1 h with TBS containing 0.2% Tween 20, 5% non-fat dry milk, incubated for 30 min at 37°C in a microplate shaker at 750 rpm with 1:100 dilution of anti-rTMPT2A in TBS, and washed five times with TBS containing 0.05% Tween 20. Bound anti-mouse antibodies were detected with FITC-conjugated rabbit anti-mouse (Dako, Denmark) diluted 1:500 in TBS, after incubation for 30 min with shaking. After five washes in TBS, 100 μL of TBS was added to each well, and the fluorescence was measured in a microplate fluorescence reader (Bio-Tek Instruments, USA), at an excitation wavelength of 490 nm, and emission wavelength of 525 nm (sensitivity, 70%). The results are expressed in arbitrary fluorescence units.

### Reverse transcriptase-quantitative polymerase chain reaction (RT-qPCR)

Aliquots of 10^6^ trophonts/mL of *P. dicentrarchi* were incubated for 10, 60 and 240 min with turbot immune serum diluted 1:50 in incomplete L-15 medium. In some experiments, ciliates were incubated for 240 min with 500 μM of dibucaine hydrochloride (Sigma-Aldrich). Total RNA was isolated from the trophonts by using the NucleoSpin RNA isolation kit (Macherey-Nagel) according to the manufacturer’s instructions. After purification of the RNA, the quality, purity and concentration were measured in a NanoDrop ND-1000 Spectrophotometer (NanoDrop Technologies, USA). The reaction mixture (25 μL) used for cDNA synthesis contained 1⋅25 μM random hexamer primers (Promega), 250 μM of each deoxynucleoside triphosphate (dNTP), 10 mM dithiothreitol (DTT), 20 U of RNase inhibitor, 2⋅5 mM MgCl_2_, 200 U of Moloney murine leukemia virus reverse transcriptase (MMLV; Promega) in 30 mM Tris and 20 mM KCl (pH 8⋅3) and 2 μg of sample RNA. PCR (for cDNA amplification) was performed with gene-specific primers forward/reverse pair for the TMPT2A gene (FTMPT2/RTMPT2) 5’-ATTTGCTTGCGTTCTCGTCT-3’ / 5’-TCATCTTCGTCTTGGGCTCT-3’; TMPT4B gene (FTMPT4/ RTMPT4) 5’-CCACGAGAGATGGGTAGAGG-3’ / 5’-AATTCAATCTGGTGGCCAAT-3’. In parallel, a qPCR was performed with *P. dicentrarchi* elongation factor 1-alpha gene (EF-1α) (GenBank accession KF952262) as a reference gene, by including the forward/reverse primer pair (FEF1A/REF1A) 5′-TCGCTCCTTCTTGCATCGTT-3′/ 5′-TCTGGCTGGGTCGTTTTTGT-3′). The Primer 3Plus program was used, with default parameters, to design and optimize the primer sets. Quantitative PCR mixtures (10 μL) contained 5 μL Kapa SYBR FAST qPCR Master Mix (2X) (Sigma-Aldrich), 300 nM of the primer pair, 1 μL of cDNA and RNase-DNase-free water. Quantitative PCR was developed at 95 °C for 5 min, followed by 40 cycles at 95 °C for 10 s and 60 °C for 30 s, ending with melting-curve analysis at 95 °C for 15 s, 55 °C for 15 s and 95 °C for 15 s. qPCRs were performed in an Eco RT-PCR system (Illumina). Relative quantification of gene expression was determined by the 2^−ΔΔCt^ method [25] applied with software conforming to minimum information for publication of RT-qPCR experiments (MIQE) guidelines [26].

### Intracellular Ca^2+^ release analysis

The release of intracellular Ca^2+^ after stimulation of ciliates with turbot immune serum was analyzed using the No-Wash, Fluo-4NW calcium assay kit (Life Technologies). The ciliates (2×10^5^) were washed twice by centrifugation with Hanks’ balanced salt solution (HBSS without Ca^2+^, Mg^2+^, and phenol red) and resuspended in assay medium (HBSS, 20 mM HEPES and 2.5 mM probenecid) to a final concentration of 1.25×10^6^ ciliates/mL. The ciliates were then incubated with 1:50 dilution of turbot immune serum in 96-well microplates at 21°C. The cell-permeable Ca^2+^ indicator probe, Fluo-4 NW, was added following the manufacturer’s instructions, and the fluorescence (Ex: 494 nm, Em: 516 nm) was measured in a fluorimeter (FLx800, BioTek, USA). Negative controls with HBSS were included.

### Bioinformatic and statistical analysis

The performance of the functional analysis of proteins was evaluated and families predicting the domains and important sites were classified using InterPro software [27]. The transmembrane topology and location of signal peptide cleavage sites in amino acid sequences were predicted using Phobius [28], SignalP [29] and Signal-3L 2.0 [30] software. Protein sequence motifs were searched for using the MotiFinder tool of the Japanese network GenomeNet (accessible on-line at: https://www.genome.jp/tools/motif/MOTIF.html). Mucin type GalNAc O-glycosylation sites were predicted using the NetOGlyc 4.0 Server [31]. The physicochemical parameters for a given protein were predicted using the ProtParam tool [32]. Protein modelling was conducted using the SWISS-MODEL server [33]. The cysteine and histidine metal binding sites of the sequenced protein were predicted using METALDETECTOR v2.0 [34]. The amino acid sequences of the TMPT2A and TMPT4B genes were aligned using Clustal Omega [35]. The evolutionary history was inferred using the Maximum Likelihood method based on the JTT matrix-based model [36]. Finally, evolutionary analyses were conducted in Mega7 [37].

The results are expressed as means ± standard error of the means (SEM). The data were examined by one-way analysis of variance (ANOVA) followed by the Tukey-Kramer test for multiple comparisons, and differences were considered significant at *α*=0.05.

## RESULTS

### Morphological changes produced in ciliates incubated with immune serum from the host

Immunization of turbot with a crude extract of ciliates-CL-generated sufficient levels of antibodies to induce immobilization/agglutination, with peak levels reached after one hour of incubation (Fig. 1). As already indicated, we used inactivated immune sera to prevent the lytic action of complement and to enable specific study of the processes produced exclusively by the action of the antibodies during agglutination/ immobilization of the trophonts.

**Figure 1.**
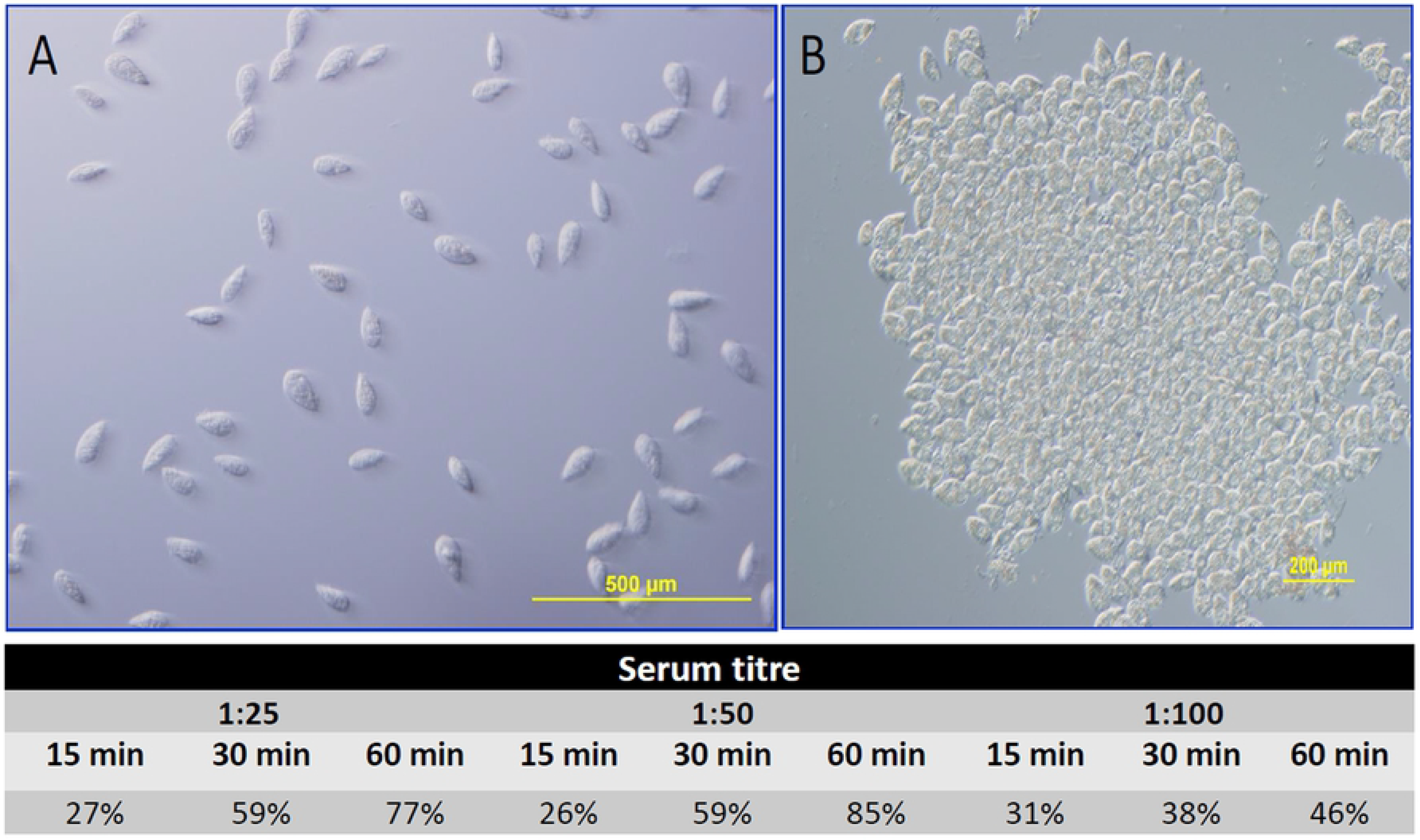
Microphotographs obtained by differential interference contrast microscopy showing *P. dicentrarchi* trophonts after being incubated for 30 min with (A) preimmune serum, and (B) immune serum. The lower table shows the effect of different dilutions of turbot immune serum (antibody titre) and different incubation times on agglutination of ciliates (results expressed as percentages).

The presence of agglutinating antibodies caused the agglutinated trophonts to produce a mucoid capsule, which became increasingly evident throughout the incubation period. After two hours of incubation, the ciliates began to emerge from the capsules, showing a normal morphology, and the number of free ciliates increased over time. The empty capsules showed the external morphology of the parasite. (Fig. 2). SEM-examination of the agglutination process clearly revealed the superficial changes that take place in the ciliate in the presence of the turbot immune serum over time (Fig. 3). The addition of the immune serum initially did not seem to affect the ciliates, whose ciliary morphology was apparently unchanged (time 0); however, during the incubation period the trophonts increased in diameter and a layer of gradually thicker amorphous material appeared on the surface. At the end of the process, microphotographs clearly show the presence of structures that maintain the external ciliary morphology but that are hollow. The free ciliates showed a normal ciliary structure (Fig. 3).

**Figure 2.**
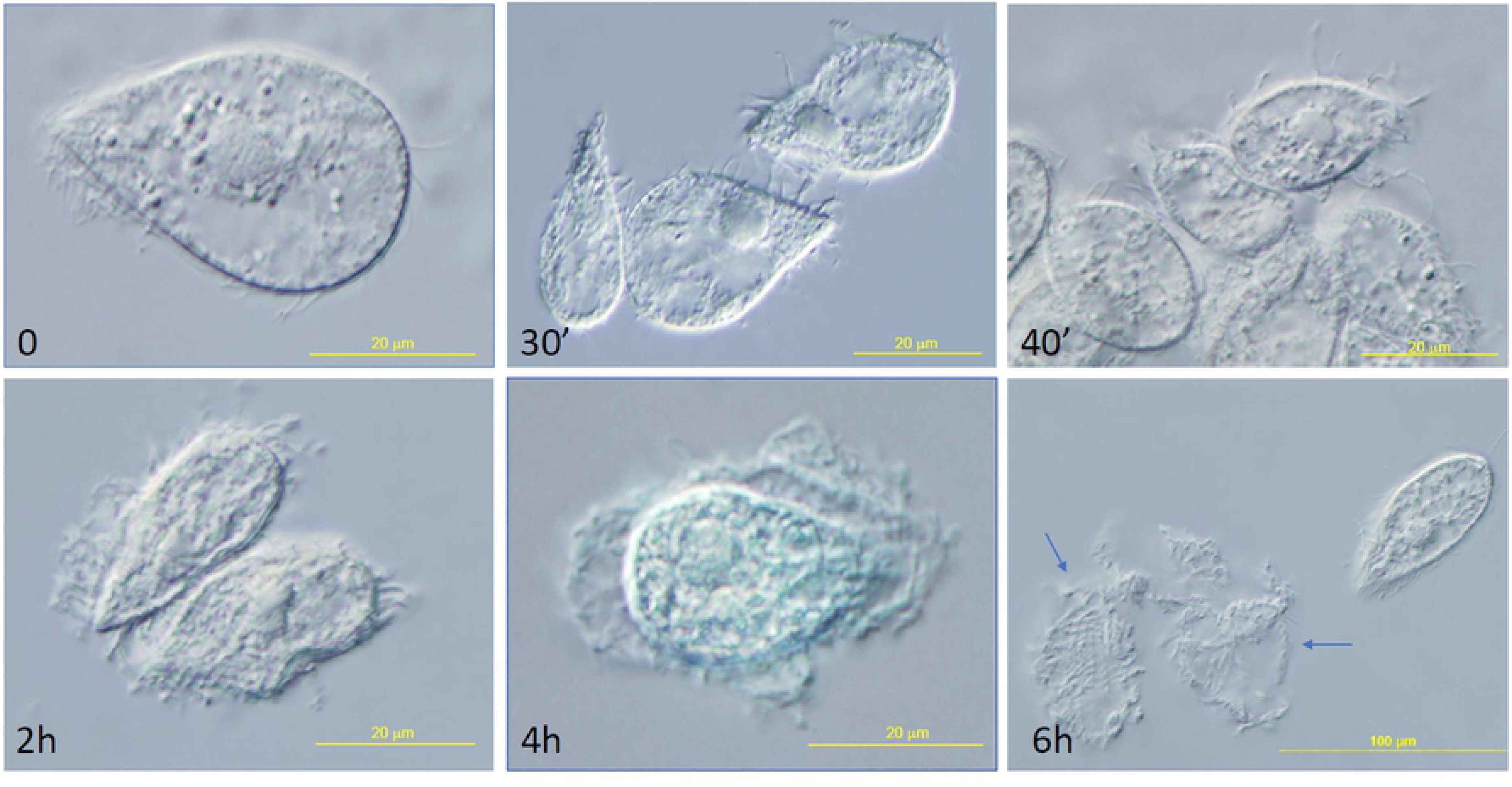
Microphotographs obtained by differential interference contrast microscopy showing the sequence of changes after agglutination of the *P. dicentrarchi* trophonts caused by the addition of the turbot immune serum (up to 6 h incubation), including the presence of empty capsules (arrows).

**Figure 3.**
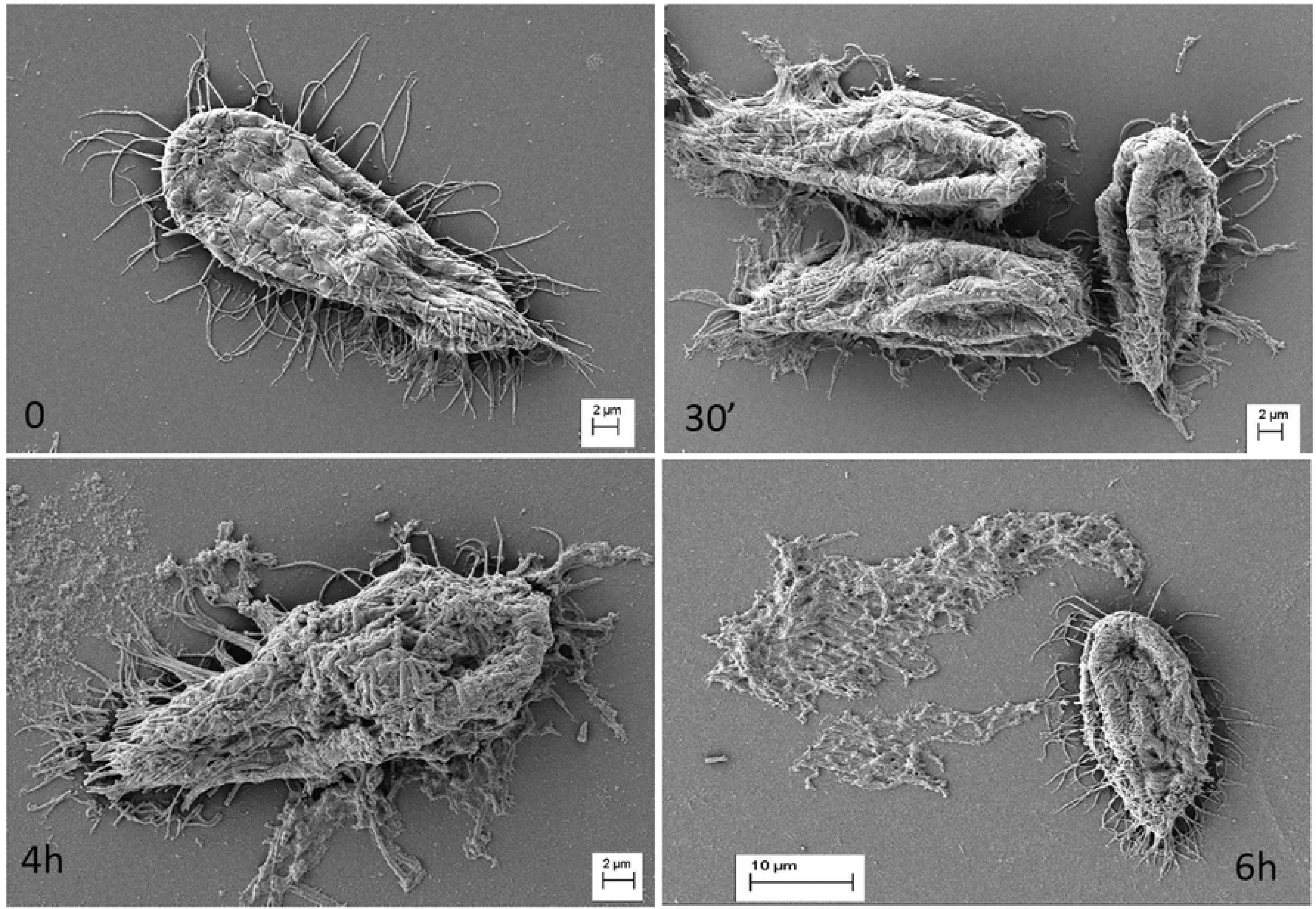
Microphotographs obtained by scanning electron microscope (SEM) of *P. dicentrarchi* trophonts, showing the changes in the ciliate surface after the addition of the turbot immune serum (up to 6 h incubation).

### Molecular and biochemical characterization of the extrusome proteins

*P. dicentrarchi* has two types of extrusomes associated with the plasma membrane and located at the insertion sites between the alveolar sacs (Fig 4). Examination by electron microscopy revealed that the extrusomes have spherical morphology (Fig. 4A) or elongated morphology (Fig. 4B). Apart from the morphology, the characteristics of the material that these two types of extrusomes contain were also different. Thus, on the one hand, rounded extrusomes contained an electrolucent material (Fig. 4A), while elongated extrusomes contained greater amounts of electrodense material (Fig. 4B).

**Figure 4.**
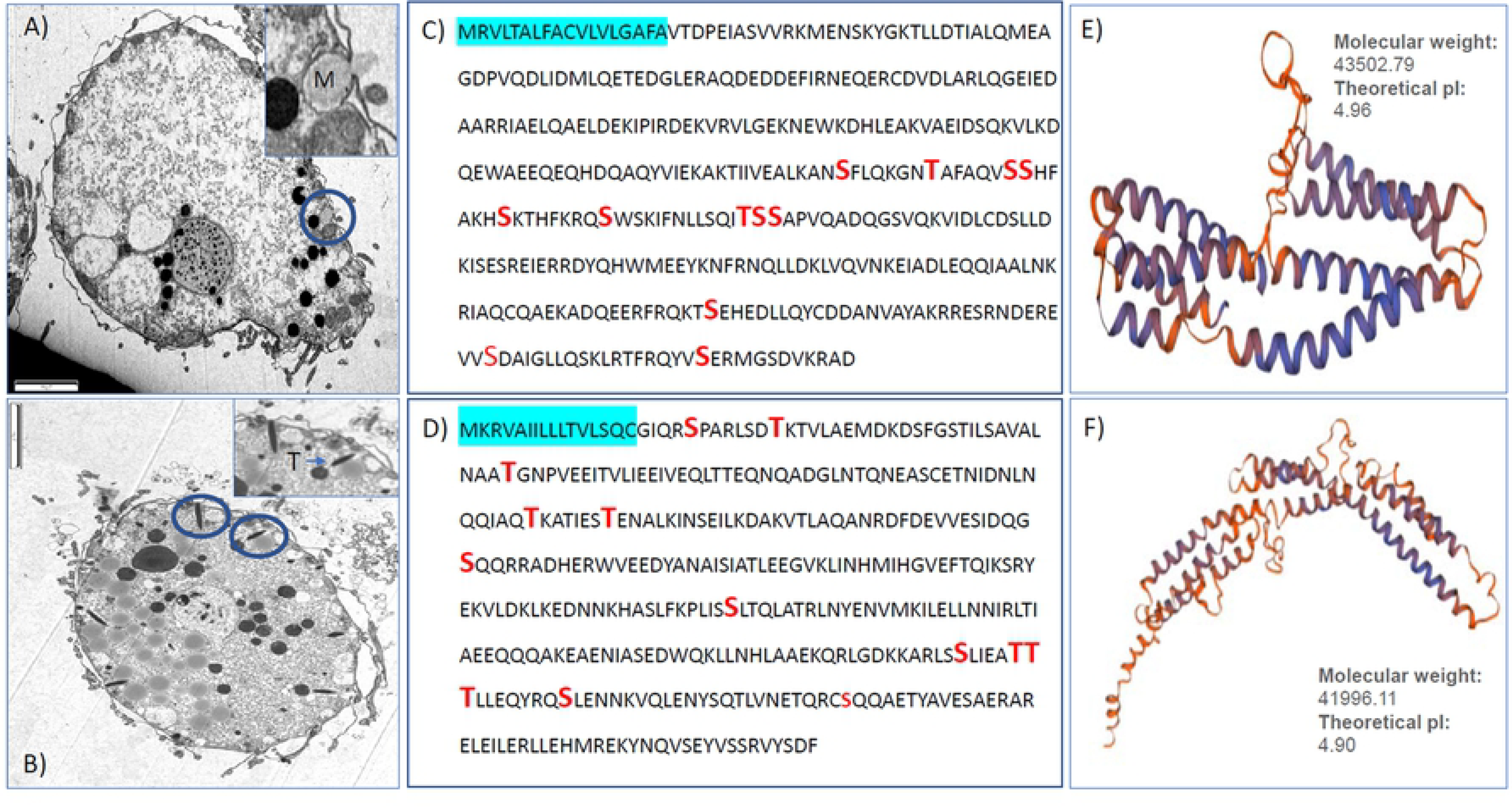
(A,B) Microphotographs obtained by transmission electron microscopy (TEM) of *P. dicentrarchi* trophonts showing the structure of the two basic types of extrusomes: A) spherical extrusomes (circle) of mucocyst type (M), and a detailed enlargement of these structures in the upper right-hand side of the image; (B) fusiform extrusomes (circle) of trichocyst type (T), and a detailed enlargement of these structures in the upper right-hand side of the Image. C-D) amino acid sequence of *P. dicentrarchi* trichocyst matrix protein T2-A and T4-B (TMPT2A and TMPT4B, respectively). The shaded region indicates the prediction of a signal peptide between aa 1-18 (C) and aa 1-16 (D); bold red indicates the potential O-glycosylation sites of the proteins. Homology modelling (Swiss-model) including molecular weight prediction and theoretical isoelectric point (ip) of the TMPT2A (E) and TMPT4B (F) proteins of the *P. dicentrarchi* trichocyst matrix. Scale bar = 2 μm.

In order to identify possible proteins contained in the extrusomes, we used RNA sequencing technology. This enabled us to sequence the entire transcriptome of the ciliates and to locate the protein sequences that may be related to the extrusomes. After annotation of the genes that encode proteins of the parasite, using the BLASTx tool, we were able to detect proteins associated with extrusomes in other ciliates. Thus, homology analysis enabled us to detect two types of proteins related to extrusomes: 1) *P. dicentrarchi* trichocyst matrix protein T2-A (TMPT2A) (accession MH412657.1) encoded by an 1134 bp mRNA that generates a protein of 377 aa (Fig. 4C), of molecular weight 43502.79 daltons and a theoretical pI of 4.96 (Fig. 4E). This protein has a signal peptide between position 1 and 18 (Fig. 4C), with a cleavage site between position 18 and 19, corresponding to aa 15-18 with the signal peptide C-region, between aa 3-14 the signal peptide H-region is located and between the aa 1-2 the signal peptide N-region. The TMPT2A protein possesses 12 potential O-glycosylation sites in the aa 182, 189, 195-196, 202, 210, 222-224, 317, 348 y 366 (Fig. 4C), and binds to metals in the cysteine at position 10. 2) *P. dicentrarchi* trichocyst matrix protein T4-B (TMPT4B) (accession MH412658.1) encoded by an 1113 bp mRNA and possesses 370 aa (Fig. 4D) of molecular weight of 41996.11 daltons and a theorical pI of 4.90 (Fig. 4F). The modeling of the structure of the proteins, using the Swiss-model, indicates that the oligomeric state of the two *P. dicentrarchi* trichocyst matrix proteins is monomeric. The protein has a signal peptide located between aa 1-16, according to the prediction by the Phobius program (Fig. 4D); however, the Signal-3L program predicts a signal peptide between aa 1 and 23. According to Phobius, the signal peptide C-region is located between aa 13 and 16, the signal peptide H-region between aa 4-12, and the signal peptide N-region between aa 1 and 3. This protein has 12 potential O-glycosylation sites in aas 21, 28, 54, 104, 111, 147, 218, 286, 291-292, 301 and 324 (Fig. 4D). BLAST analysis of the database including the *Tetrahymena thermophila* genome (TGD) indicated that this protein is related to a similar protein encoded by the GRL3 gene (Granule Lattice), which encodes the granule lattice protein and corresponds to an acidic, calcium-binding structural protein of dense core granules, contains coiled-coil region. This protein seems to possess a cysteine at position 16 (which may be a Ca^2+^binding site), according to the prediction with METALDECTETOR v2.0 (cysteine and histidine metal binding sites predictor).

The TMPT2A and TMPT4B proteins displayed very low sequence identity (23%). The sequence identity was also very low in comparison with other ciliated proteins (e.g. *Paramecium, Ichtthyophthirius* and *Tetrahymena*) with maximum sequence identity scarcely exceeding 30%, for TMPT2A and TMPT4B (Figs. 5A, B; 6A, B). Phylogenetically, the TMPT2A protein is closer to *Paramecium* (Fig. 5C), while the TMPT4B protein is more phylogenetically spaced with the other ciliates analyzed (Fig. 6C).

**Figure 5.**
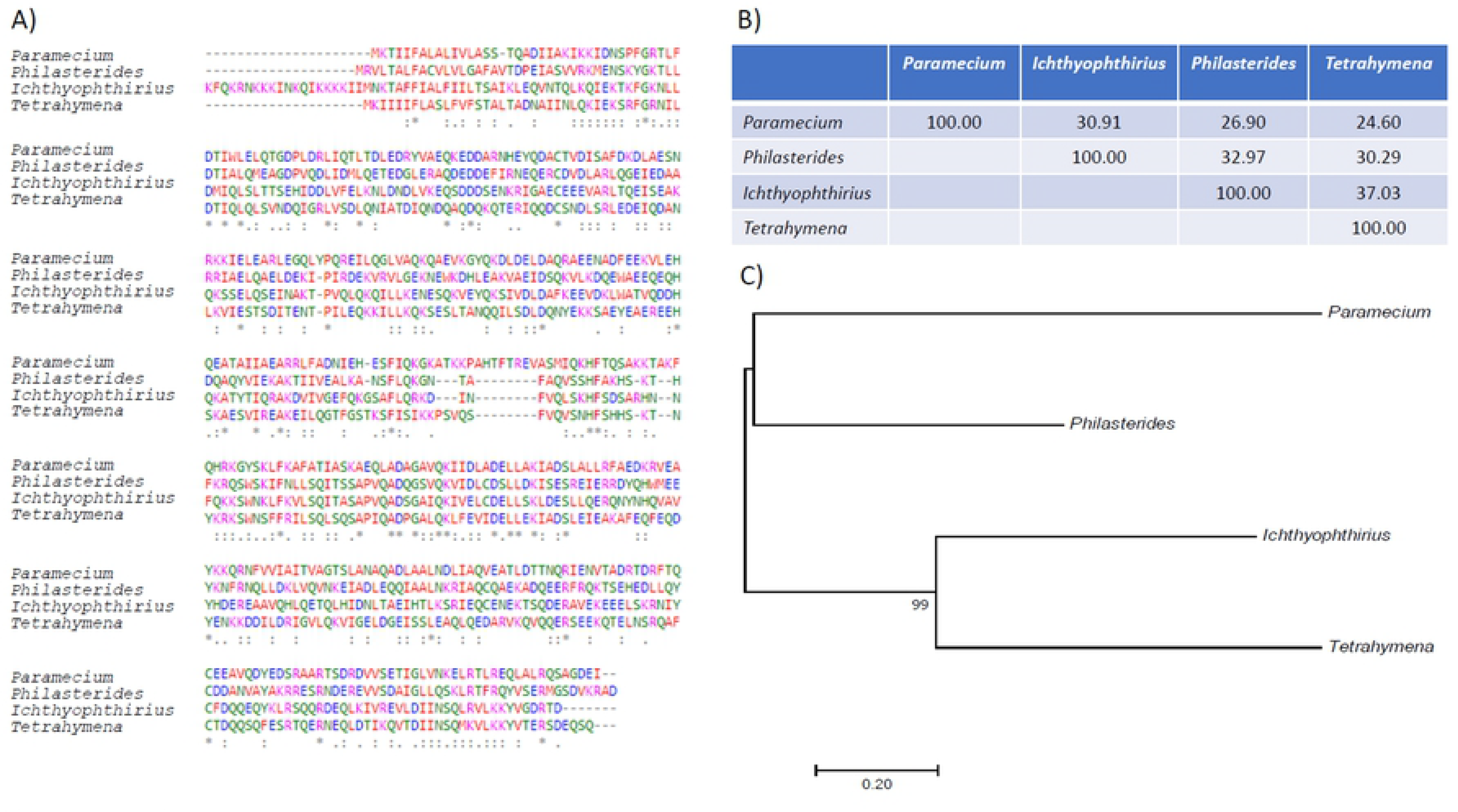
A) CLUSTAL OMEGA (v.1.2.4) multiple sequence alignment of *P. dicentrarchi* trichocyst matrix protein T2-A from four representative ciliates of the phylum Ciliophora. B) Percent Identity Matrix -created by Clustal 2.1 C) Molecular Phylogenetic analysis by Maximum Likelihood method. The tree is drawn to scale, with branch lengths measured in the number of substitutions per site. The analysis involved 4 amino acid sequences. All positions containing gaps and missing data were eliminated. The final data set included a total of 367 positions. Evolutionary analysis was conducted with MEGA7 software.

**Figure 6.**
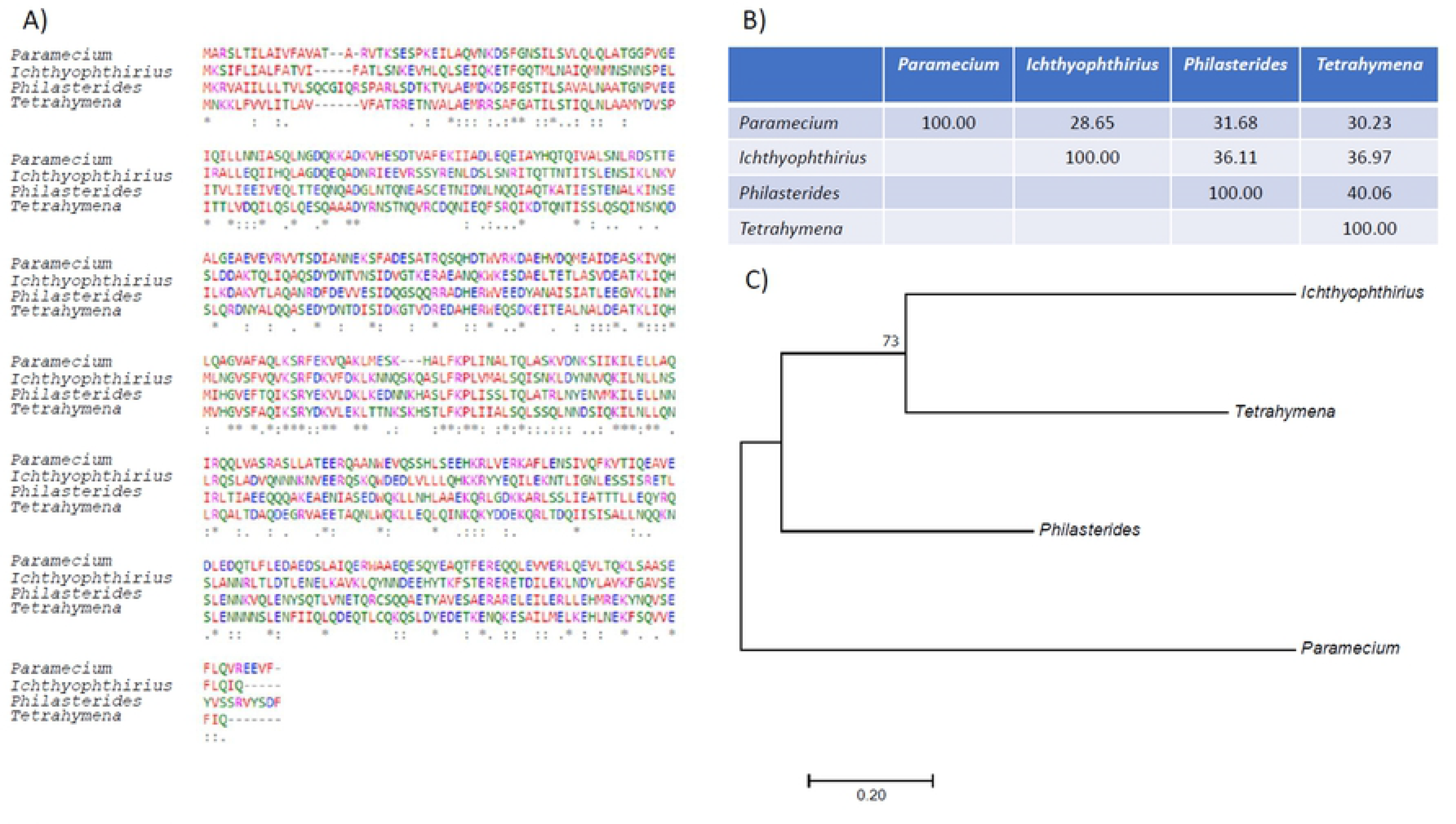
A) CLUSTAL OMEGA (1.2.4) multiple sequence alignment of *P. dicentrarchi* trichocyst matrix protein T4-B of four representative ciliates of the phylum Ciliophora B) Percent Identity Matrix - created by Clustal 2.1 C) Molecular Phylogenetic analysis by Maximum Likelihood method. The tree is drawn to scale, with branch lengths measured in the number of substitutions per site. The analysis involved 4 amino acid sequences. All positions containing gaps and missing data were eliminated. The data set included a total of 354 positions. Evolutionary analysis was conducted with MEGA7 software.

When the ciliates incubated with turbot immune serum were stained with safranin-O dye, a progressive and time-dependent increase in the intensity of staining both in the cytoplasm and in the external material surrounding the ciliate was observed (Fig. 7).

**Figure 7.**
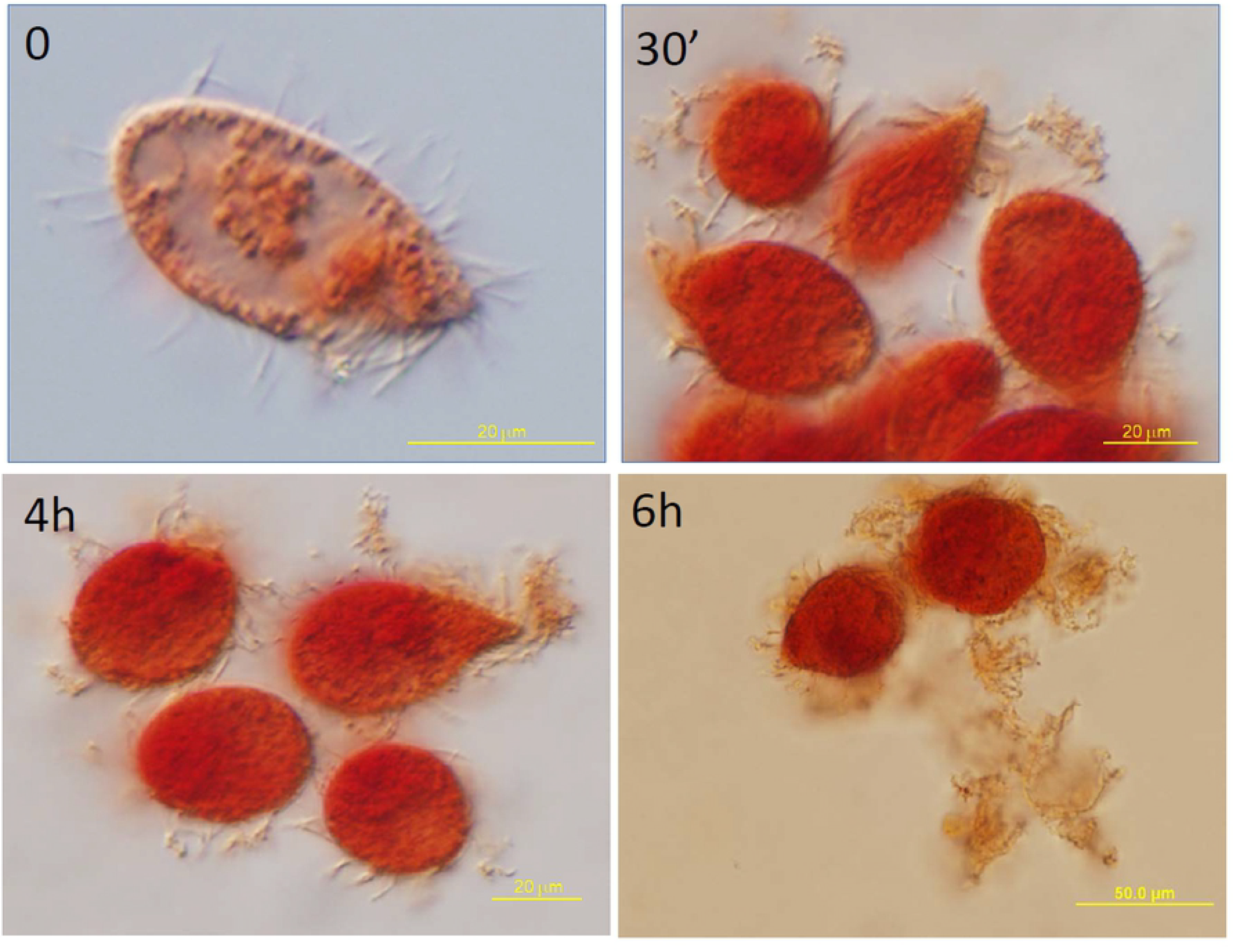
Histochemistry analysis of mucin production (a peptidoglycan component of extrusomes) by safranin O staining of *P. dicentrarchi* trophonts after incubation with A) control serum, B) immune turbot serum for 30 min, C) immune turbot serum for 2 h, D) with immune turbot serum for 6h.

### Expression and location of extrusome proteins after stimulation with immune serum from the host

In order to determine whether the proteins presumably associated with the trichocysts are involved in the formation of the capsules observed during the agglutination of the ciliates by the host immune serum, the recombinant protein was generated in the yeast *Kluyveromyces lactis*. For this purpose, we expressed the TMPT2A protein in the yeast (Fig. 8A), which was used to generate antisera in mice to enable us to perform experiments to study expression of this protein after incubation with the antiserum (Fig. 8B) and to determine the cytolocation (Fig. 8C).

**Figure 8.**
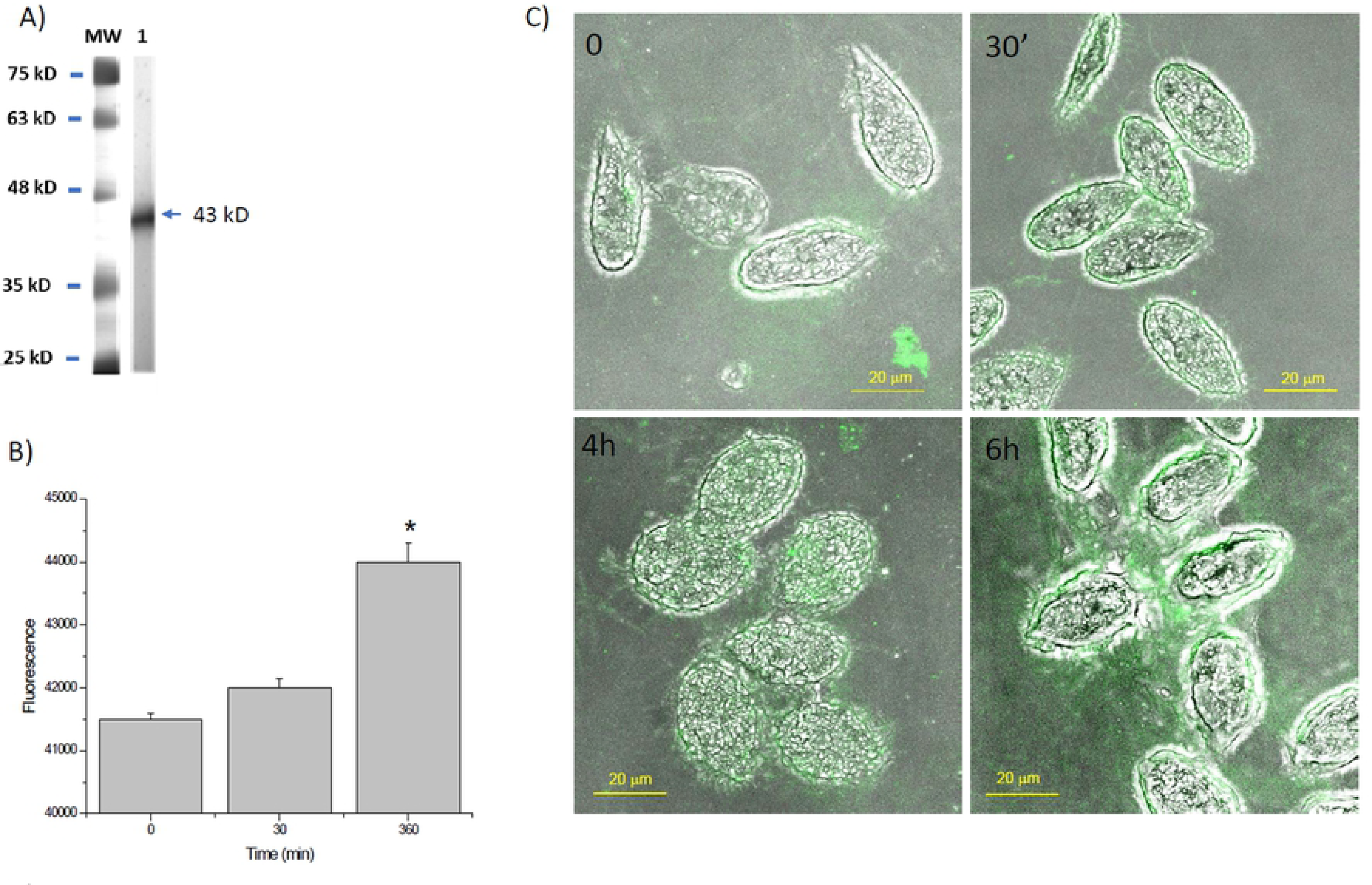
(A) SDS-PAGE analysis of the recombinant *P. dicentrarchi* trichocyst matrix protein T2-A (rTMPT2A), lane 1. MW: Molecular weight markers in kD. (B) FELISA of levels of TMPT2A expressed by the trophonts incubated for 30 min and 6 h with the turbot immune serum. Values are means ± standard errors. The symbol indicates a significant difference (*P <* 0.01) relative to the control (time 0). (C) Microphotographs obtained by confocal / phase contrast microscopy of *P. dicentrarchi* trophonts incubated with immune serum for different times. The images correspond to the combination of a visible image and an immunofluorescence (green signal) using a recombinant mouse antibody anti-*P. dicentrarchi* TMPT2A and revealed with an anti-mouse rabbit antibody conjugated with FITC.

First, the recombinant protein expressed by yeast has the biochemical characteristics (e.g. molecular size) predicted for the original sequence obtained from the ciliate, which indicates that this protein expression system is optimal for the heterologous expression of this type of eukaryotic proteins (Fig. 8A). On the other hand, the antibodies generated in mice against the rTMPT2A protein demonstrated that the material produced after incubation of ciliates with the turbot immune serum is related to this protein, as demonstrated by the FELISA assay, in which the absorbance levels of these antibodies increase during the period of incubation with the immune serum from turbot (Fig. 8B). By using immunofluorescence, an increase in fluorescence was observed in both the cytoplasm of the agglutinated ciliates and in the material associated with the outer surface throughout the incubation period (Fig. 8C).

#### Expression of the genes associated with extrusome proteins and their association with the discharge of intracellular Ca^2+^ after stimulation of the ciliates with host immune serum

We investigated expression of the genes encoding the trichocysts proteins TMPT2A, TMPT4B after incubation with the turbot immune serum for different times. Incubation of the ciliates with the turbot immune serum produced a significant increase in the mRNA levels of the genes encoding these proteins throughout the incubation period (Fig. 9A). Dibucaine, included as a positive control for the induction of extrusion, also had a stimulatory effect on the expression of both mRNA levels relative to the all trichocyst genes; however, the absolute mean values of increase were higher for the TMPT2A gene than for the TMPT4B gene (Fig. 9A).

**Figure 9.**
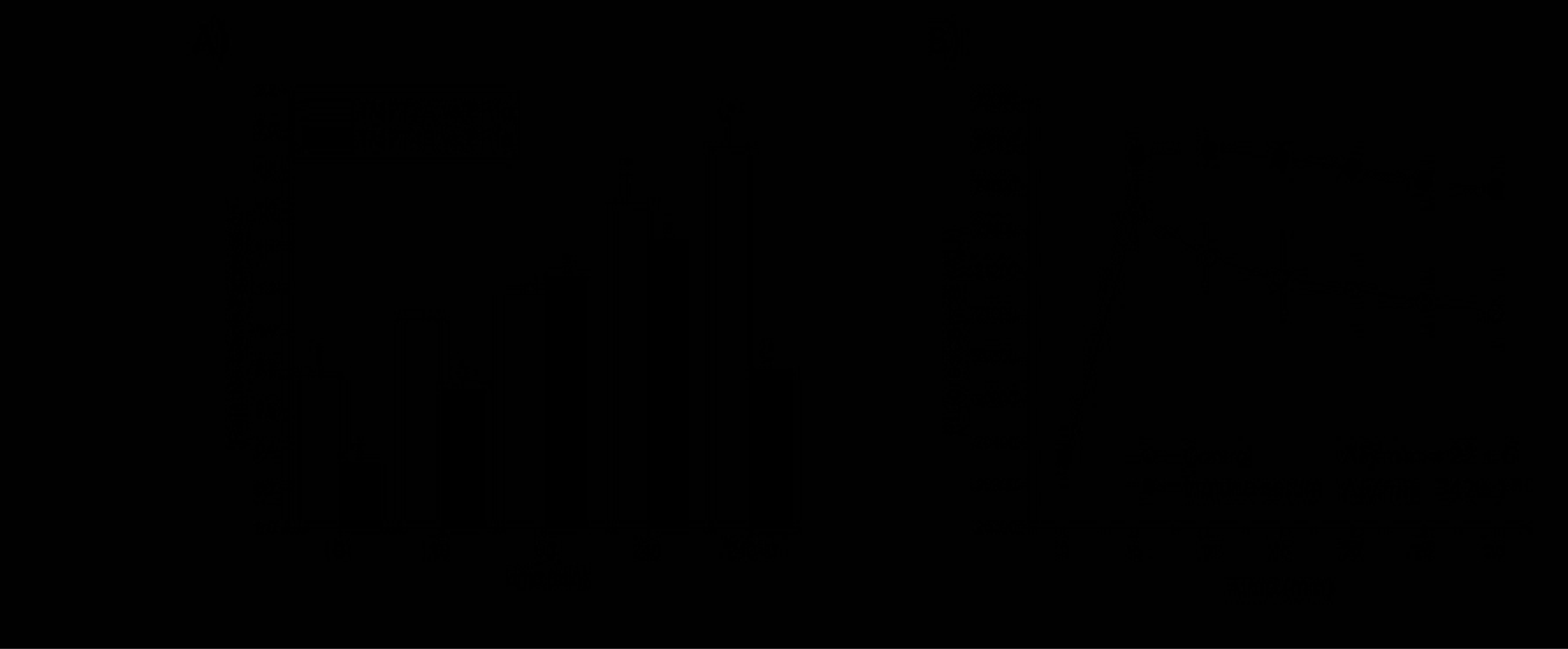
(A) Levels of mRNA expression of the genes that encode the *P. dicentrarchi* trichocyst matrix protein T2-A (TMPT2A) and *P. dicentrarchi* trichocyst matrix protein T4-B (TMPT4B) in ciliates incubated for different lengths of time with turbot immune serum and dibucaine (D). The results are expressed as the relative gene expression versus the *P. dicentrarchi* elongation factor 1-alpha (EF1α). (B) Calcium response of trophonts stimulated with turbot immune serum (ITS) and Hanks’ balanced salt solution (HBSS without Ca^2+^, Mg^2+^, and phenol red) quantified using the Fluo-4 NW calcium assay kit. The time course of the increase in fluorescence by min (ΔF/min) of cell-permeable fluorescent dye reflects the rates of dye-loading of cells by passive uptake of the AM esters and the influx of calcium through membrane channels or release from intercellular stores. The values at each data point are the mean ± standard error (SE) for five replicates. Asterisks indicate a statistically significant difference (*P*<0.01) relative to control (time 0).

Finally, we analyzed the effect of the addition of turbot immune serum on the intracellular Ca^2+^ discharge by using the Fluo-4NW probe. Incubation of the trophonts with the turbot immune serum induced discharge of intracellular Ca^2+^ discharge, as evidenced by the increase in fluorescence levels throughout the incubation time, while the fluorescence increased only slightly over time in the ciliates not exposed to the serum (Fig. 9B).

## Discussion

In protists, extrusomes are specialized exocytotic and ejectable organelles which can discharge their contents outside of the cell in response to external mechanical or chemical stimuli and which can have offensive or defensive functions during predation or in the acquisition of food [38]. In *P. dicentrarchi*, two types of extrusomes have been described: a fusiform type (fibrous trichocysts) located in the cortex, perpendicular to the plasma membrane, and a spherical type (mucocysts) with an irregular distribution [18,19]. The mucocysts, which have an amorphous content, merge with the plasma membrane and release their contents to the exterior giving rise to a thin mucilaginous layer over the cell surface [19]. Although in free-living ciliates the extrusomes can have a protective or defensive response to environmental changes, in ciliated parasites such as *P. dicentrarchi*, the extrusomes may play a role in providing protection from attack by the host immune system. The existence of the production of capsules by the trophonts of *P. dicentrarchi* was initially obtained in studies of ciliate agglutination caused by different immune sera from turbot and rabbit [20]. In those studies, it was observed that when ciliates were incubated with the immune sera (for 2h), abundant transparent capsule-like structures appeared. The precise surface topography of the ciliate, including the somatic cilia could be seen and ciliates were also observed moving within the capsules [20]. At that time, it was interpreted that their capsules probably made up of immunocomplexes between these antigens and the agglutinating antibodies [20]. In the present study, we sequentially monitored the agglutination of the trophonts by inactivated immune turbot serum in order to investigate the capsule formation. The phenomenon of capsule formation has already been described in the ciliates; e.g. *Tetrahymena* forms capsules when exocytosis of mature mucocysts is induced by the secretagogue Alcian Blue 8GS [39–41]. In the environment, the ciliate mucocysts secrete an amorphous material to protect the cell from osmotic shock or from predator attacks [43].

The appearance of capsules during agglutination of the *P. dicentrarchi* trophonts with immune serum suggests that the host antibodies induce the mucocysts to extrude their mucilaginous content. This material is deposited on the surface of the ciliate forming a protective layer, which eventually became a rigid capsule with an external topology identical to that of the ciliate and which protects it from agglutination. This process was clearly observed in this study by both optical microscopy and SEM.

In ciliates such as *Paramecium*, trichocysts are characterized by a highly constrained shape that reflects the crystalline organization of the proteins that they contain and that are derived from the process of a broad family of precursor proteins (coded by a family of some 100 coexpressed genes) that allow correct processing of the crystalline core assembly necessary for functioning of the trichocyst [44,45]. The trichocyst matrix proteins in *Paramecium* are of sizes ranging between 15-20 kDa, and some are glycosylated; the isoelectric points are between 4.7 and 5.5 and the proteins seem to be derived from the proteolytic processing of precursor proteins of size between 40-45 kDa [46,47]. In our study, the TMPT2A and TMPT4B proteins were about 43 kDa in size and the isoelectric points were close to 5.0, i.e. they are compatible with the precursor proteins described in *Paramecium.* In addition, the proteins from *P. dicentrarchi* possess sequences with a very low similarity to each other, although with very similar isoelectric points and sizes. This may indicate that the trichocyst matrix is composed of complex interrelated proteins, or of the proteolytic processing during the maturation of secretory proteins [46], or of post-translational modifications [48]. It has also been observed in *Paramecium tetraurelia* that proteins released by exocytosis of trichocysts are glycoproteins [49].

As previously mentioned, apart from the encysting stages of the ciliates, capsule production is rare, but has been induced *in vitro* in several species [42]. The capsule has been shown to consist of mucopolysaccharide material from mucocysts [50,51]. *Tetrahymena* has mucocyst-type extrusomes characterized by containing mucin-like acidic proteins of sizes between 40 and 80 kDa and that can bind to Ca^2+^ [8]. O-glycosylation (or “mucin-type O-glycosylation”) indicates that these proteins carry this type of glycan to the side-arm hydroxyl groups of serine and threonine residues [52]. Safranin O staining has been used to detect glycosaminoglycans [53] and mucins [54]. All mucins are highly O-glycosylated, and the biosynthesis and degradation are perfectly integrated for protection of the cell against external aggressions [55]. The present findings clearly show that the presence of antibodies in the turbot immune serum acts as a stimulus that leads to the production of mucin-like proteins, as shown by Safranin staining. The stimulation also causes a significant increase in the expression of both the matrix proteins and the expression of the genes that encode them. The immunological assays revealed that the components of the capsule share epitopes with the matrix glycoproteins of the extrusomes.

In ciliate secretion systems, Ca^+2^ is necessary for stimulus-secretion coupling [56]. In *Paramecium* it has been shown that the exocytic release of the paracrystalline secretor product derived from the trichocyte matrix depends on Ca^2+^, and the secretory signal probably involves an influx of calcium [57,58]. The role of calcium in exocytosis has been demonstrated in *Paramecium* following the application of Ca^+2^ ionophores, and direct microinjection of Ca^+2^ in the cells induces exocytosis of the trichocysts [59]. On the other hand, in *Tetrahymena*, the addition of the anaesthetic dibucaine induces the synchronous secretion of mature mucocysts [60] via an increase in intracellular Ca^+2^ [61] and the release of flocculent mucin [8]. In this study, we demonstrated that stimulation of *P. dicentrarchi* trophonts with antibodies in turbot serum induces discharge of intracellular Ca^+2^ and extrusion.

In conclusion, our findings indicate that *P. dicentrarchi* can overcome the agglutination generated by the specific antibodies produced by the host by generating capsules through the extrusome-mediated secretion of O-glycosylated matrix proteins that possess mucin-like characteristics, and whose release is regulated through Ca^+2^-mediated signalling. The findings show that the ciliate uses exocytosis as a defence mechanism that probably allows evasion of the host immune response. Likewise, analysis of the extrusome matrix proteins in yeast by heterologous production technology, which has the advantage of producing glycosylated proteins, will allow us to develop recombinant proteins of potential use in vaccines for the immunoprophylaxis of scuticociliatosis in turbot.

## Acknowledgements

This study was financially supported by grants from the Ministerio de Economía y Competitividad (Spain) and Fondo Europeo de Desarrollo Regional -FEDER-(European Union) (AGL2017-83577-R) and from the Xunta de Galicia (Spain) (ED431C2017/31) and also by the PARAFISHCONTROL project, which received funding from the European Union’s Horizon 2020 research and innovation programme under grant agreement No. 634429. This publication reflects the views of the authors, and the European Commission cannot be held responsible for any use which may be made of the information contained herein

